## Supplementary figures and images for "Touch receptor end-organ innervation and function requires sensory neuron expression of the transcription factor Meis2"

### Figure 1 Supplementary 1

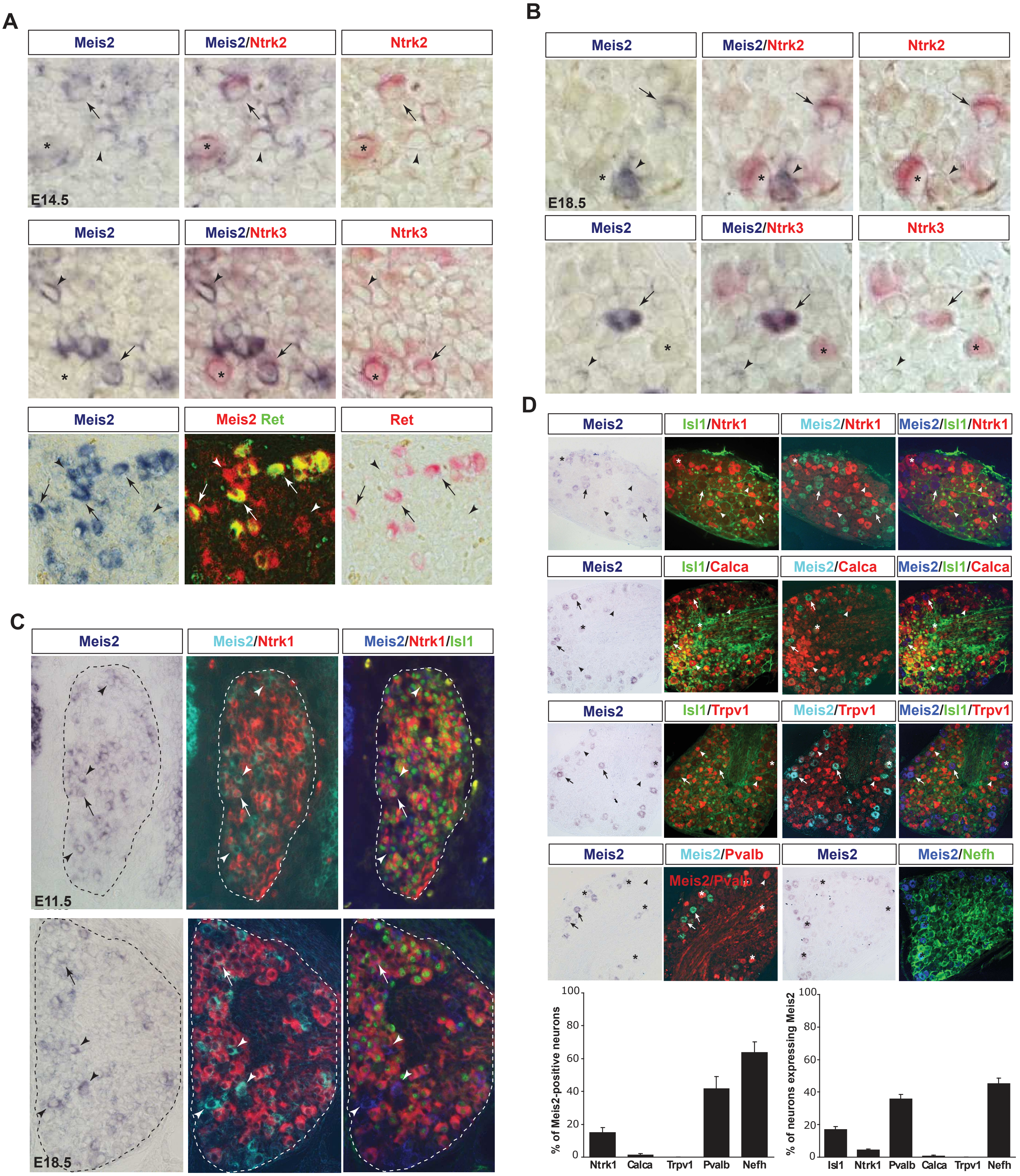


**Figure 1-Supplementary 1**

### Figure 1 Supplementary 2

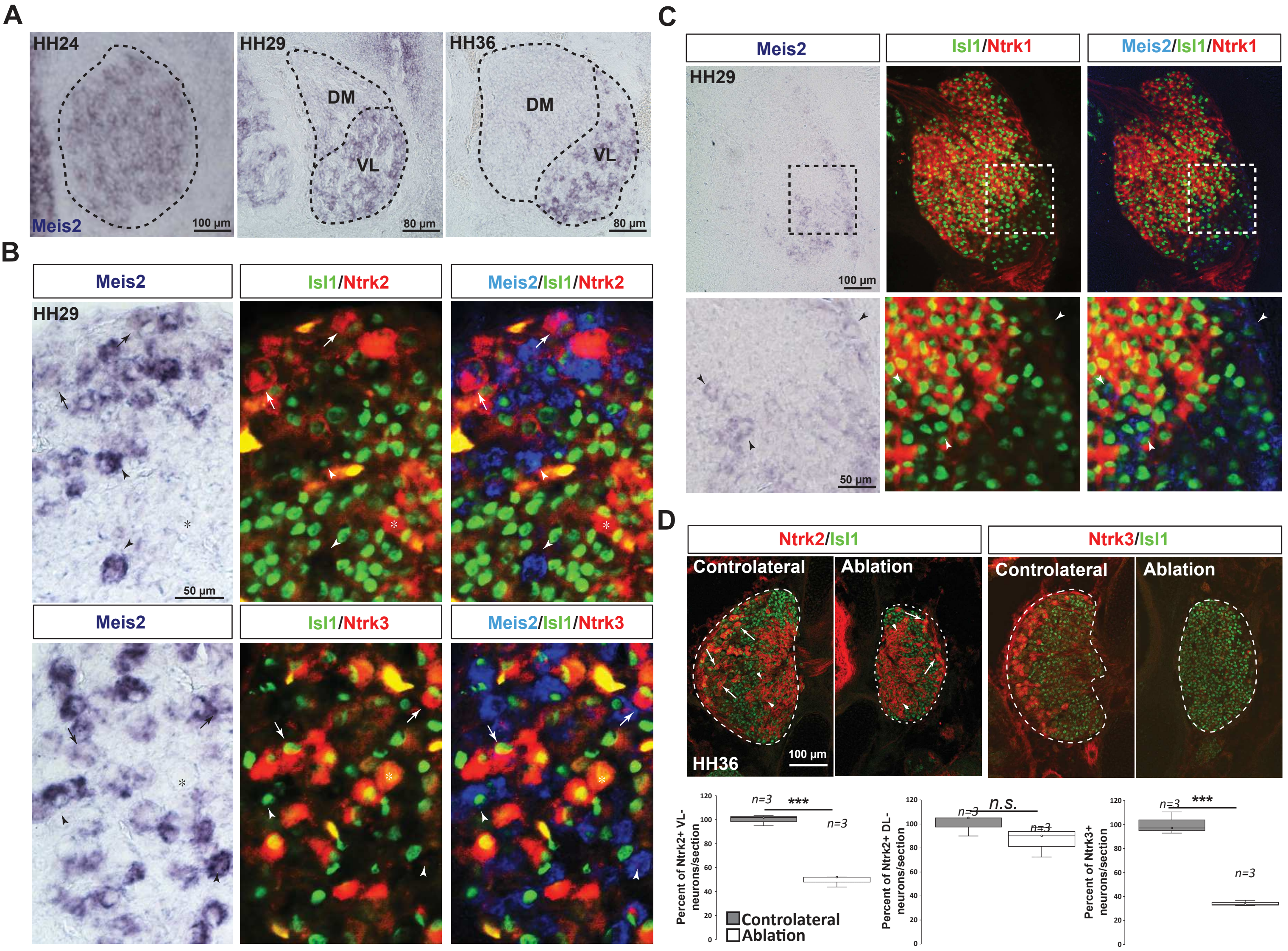


**Figure 1-Supplementary 2**

### Figure 1 Supplementary 3

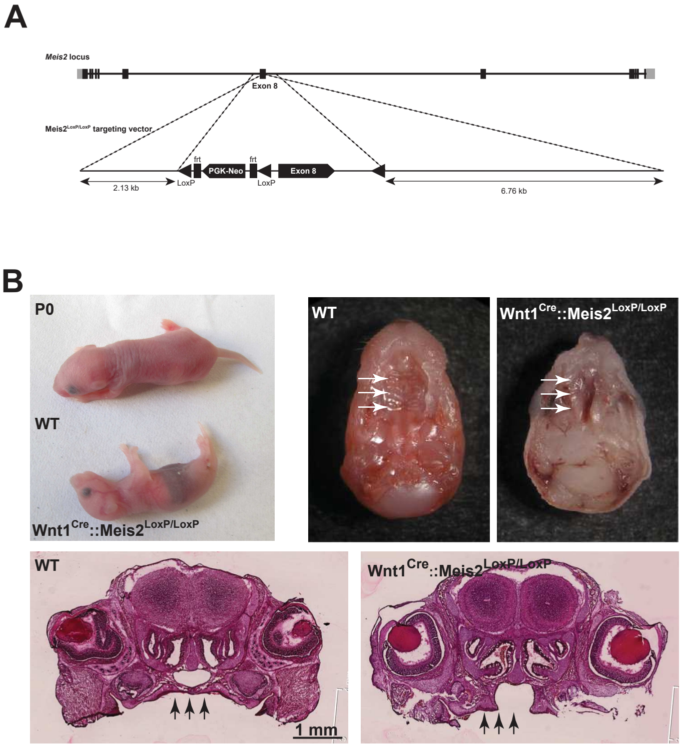


#### Figure 1-Supplementary 3

### Figure 2 Supplementary 1

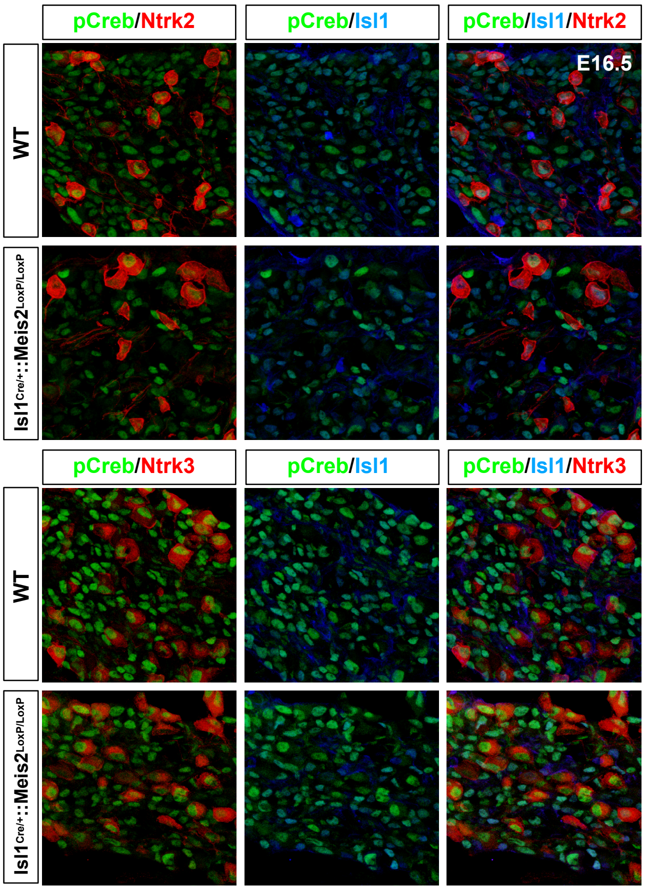


**Figure 2-Supplementary 1**

### Figure 3 Supplementary 1

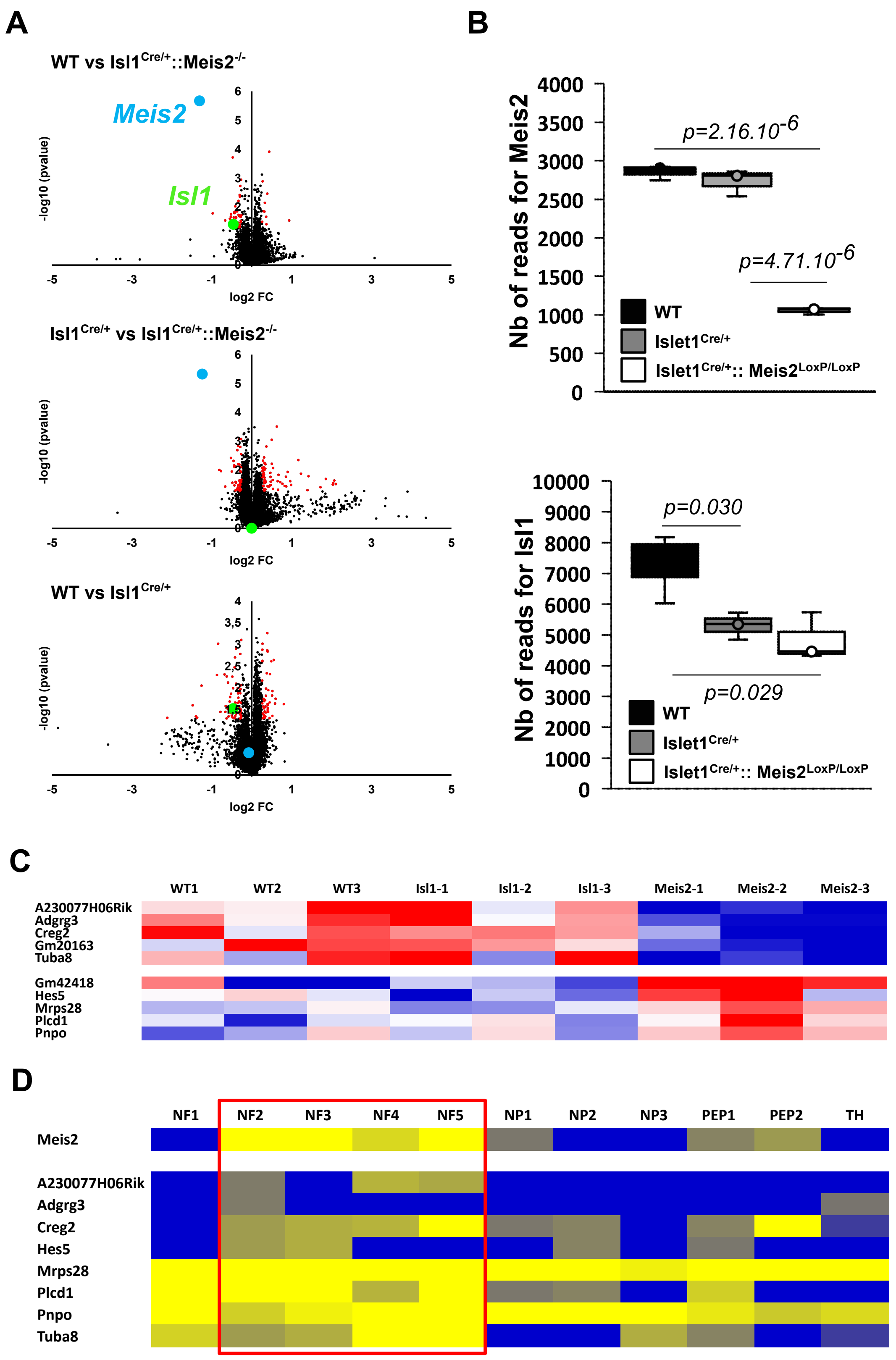


**Figure 3 Supplementary 1**

### Figure 3 Supplementary 2

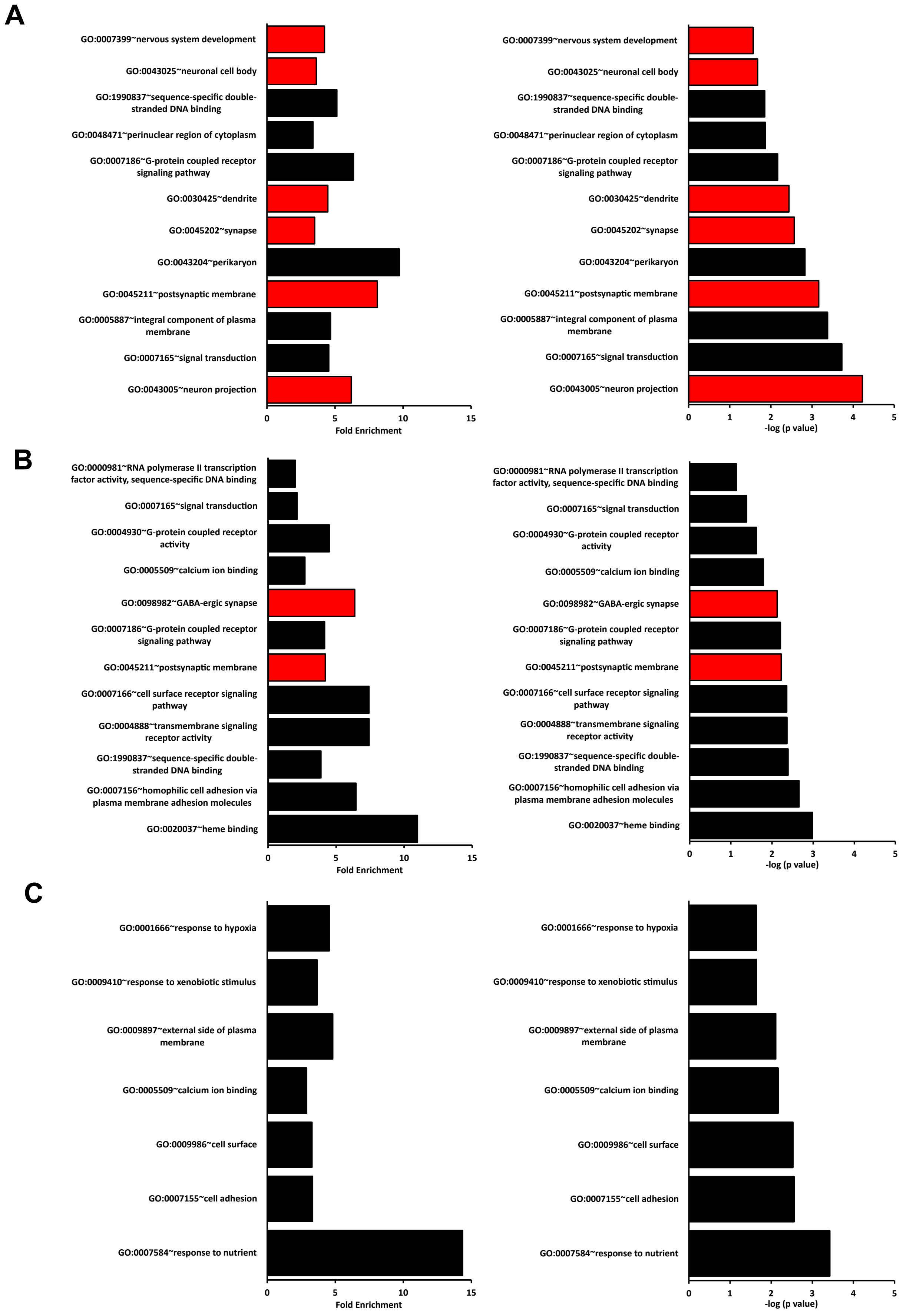


**Figure 3 Supplementary 2**

### Figure 3 Supplementary 3

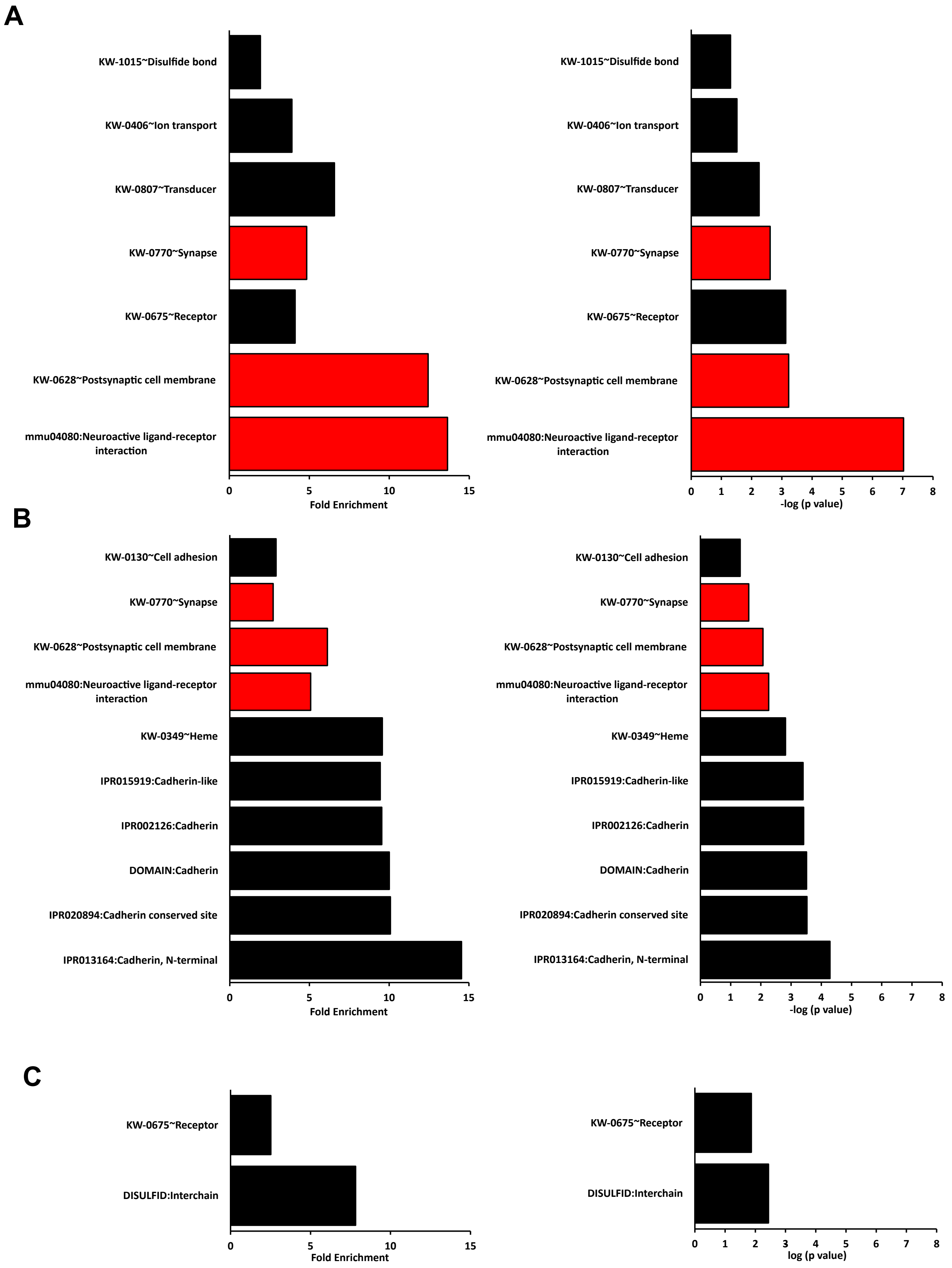


**Figure 3 Supplementary 3**

### Figure 3 Supplementary 4

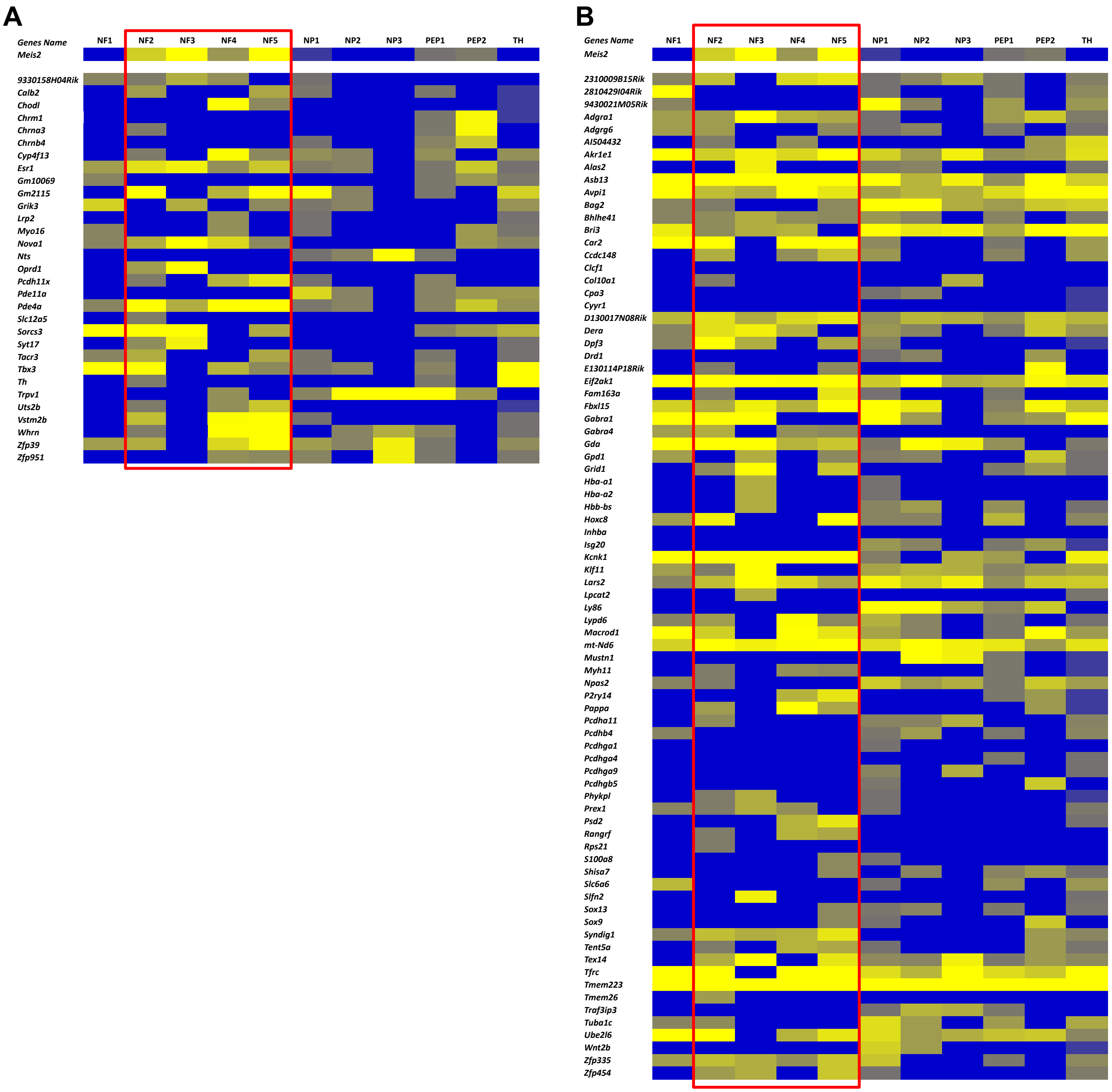

### Figure 5 Supplementary 1

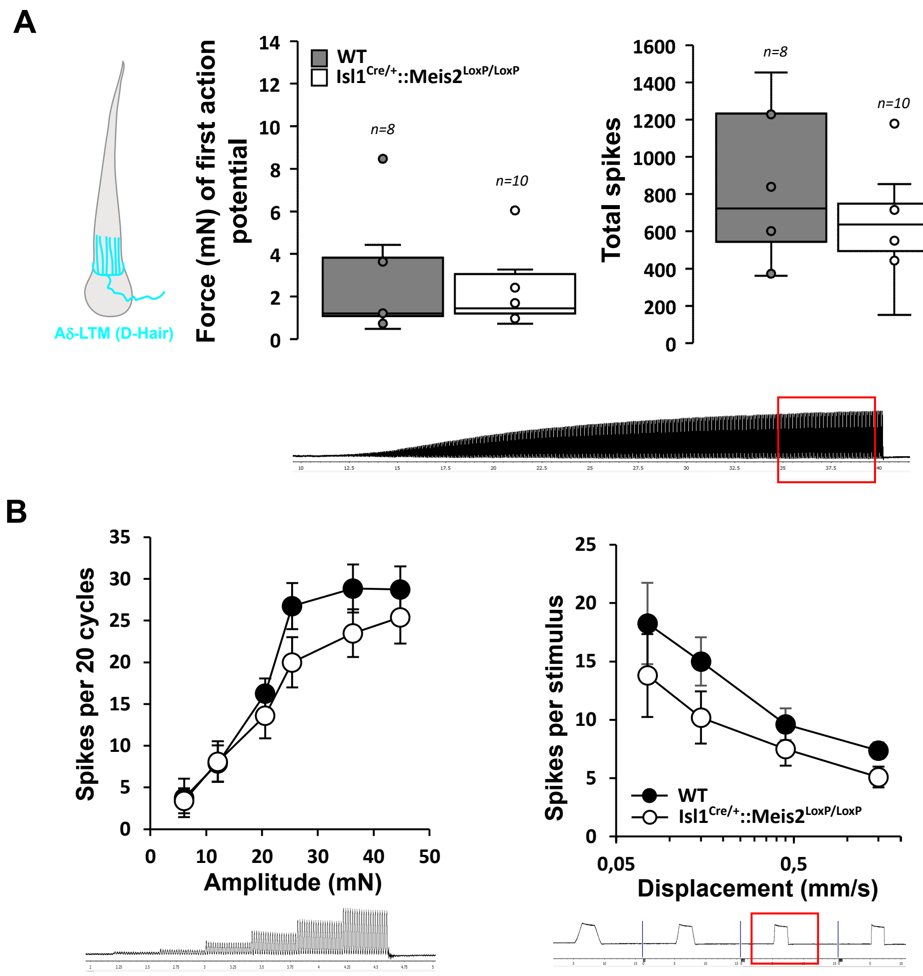


**Figure 5-Supplementary 1**
