## Supplementary material for "Touch receptor end-organ innervation and function requires sensory neuron expression of the transcription factor Meis2": Figure 1 Supplementary Table 1

Figure 1 Supplementary Table 1 : Cat walk analysis

| | **WT (n=6)** | | **Ilset1$^{Cre/+}$Meis2$^{LoxP/LoxP}$ (n=7)** | | ***t-test*** | ***t-test*** |
|---|---|---|---|---|---|---|
| **Limb** | Forelimb | Hindlimb | Forelimb | Hindlimb | *Forelimb* | *Hindlimb* |
| **Average speed (cm.s$^{-1}$)** | 39.49±4.42 | | 39.44±3.72 | | | |
| **Stand (s)** | 0,112±0,017 | 0,103±0,018 | 0,104±0,015 | 0,098±0,014 | *0,680* | *0,836* |
| **Stand Index** | -9,736±1,967 | -18,143±2,613 | -9,972±0,509 | -18,352±0,742 | *0,863* | *0,788* |
| **Max Contact At (%)** | 35,6±1,7 | 30,1±1,3 | 39,3±2,4 | 30,1±2,6 | *0,227* | *0,828* |
| **Max Contact Area (cm²)** | 0,352±0,031 | 0,352±0,011 | 0,364±0,026 | 0,370±0,028 | *0,937* | *0,397* |
| **Max Contact Max Intensity** | 218,0±0,6 | 226,1±0,5 | 219,0±1,2 | 226,7±1,6 | *0,842* | *0,865* |
| **Max Contact Mean Intensity** | 156,9±0,5 | 164,0±0,6 | 156,2±0,8 | 163,8±2,6 | *0,382* | *0,821* |
| **Print Length (cm)** | 0,962±0,033 | 0,857±0,018 | 1,002±0,044 | 0,971±0,054 | *0,670* | *0,060* |
| **Print Width (cm)** | 0,763±0,023 | 0,744±0,019 | 0,763±0,017 | 0,757±0,026 | *0,965* | *0,622* |
| **Print Area (cm²)** | 0,440±0,033 | 0,398±0,013 | 0,468±0,035 | 0,441±0,039 | *0,710* | *0,235* |
| **Max Intensity At (%)** | 17,0±2,9 | 65,2±2,8 | 23,0±1,1 | 67,8±4,2 | *0,226* | *0,603* |
| **Max Intensity** | 225,8±1,4 | 237,2±1,0 | 229,5±2,0 | 238,1±1,7 | *0,136* | *0,858* |
| **Min Intensity** | 103,0±0,9 | 105,4±1,0 | 105,0±0,8 | 107,0±1,2 | *0,139* | *0,241* |
| **Mean Intensity** | 165,8±1,1 | 173,6±1,3 | 166,0±1,5 | 172,9±2,9 | *0,755* | *0,929* |
| **Swing (s)** | 0,102±0,009 | 0,115±0,009 | 0,098±0,004 | 0,106±0,003 | *0,744* | *0,528* |
| **Swing Speed (cm/s)** | 81,0±8,3 | 71,7±7,7 | 89,2±5,0 | 81,9±5,0 | *0,632* | *0,473* |
| **Stride Length (cm)** | 7,79±0,41 | 7,80±0,38 | 8,45±0,40 | 8,45±0,41 | *0,435* | *0,439* |
| **Step Cycle (s)** | 0,213±0,026 | 0,214±0,027 | 0,194±0,007 | 0,195±0,007 | *0,661* | *0,662* |
| **Duty Cycle (%)** | 50,96±1,96 | 44,74±2,23 | 49,54±1,32 | 45,20±0,94 | *0,764* | *0,717* |
| **Toe Spread (cm)** | 0,558±0,029 | 0,632±0,017 | 0,507±0,023 | 0,626±0,032 | *0,198* | *0,924* |
| **Intermediate Toe Spread (cm)** | 0,430±0,050 | 0,398±0,020 | 0,550±0,013 | 0,415±0,023 | *0,114* | *0,995* |
| **Manual Print Length (cm)** | 0,822±0,034 | 0,729±0,026 | 0,845±0,045 | 0,832±0,055 | *0,888* | *0,093* |
| **Paw Angle Body Axis (°)** | 3,574±2,155 | 7,471±1,720 | 1,140±0,109 | 7,274±2,076 | *0,399* | *0,727* |
| **Paw Angle Movement Vector** | 5,182±2,341 | 4,920±1,221 | 5,016±1,147 | 3,735±1,587 | *0,991* | *0,960* |
| **Single Stance (s)** | 0,095±0,012 | 0,093±0,014 | 0,090±0,003 | 0,088±0,003 | *0,818* | *0,967* |
| **Initial Dual Stance (s)** | 0,008±0,003 | 0,005±0,003 | 0,004±0,001 | 0,00±0,001 | *0,372* | *0,365* |
| **Terminal Dual Stance (s)** | 0,009±0,003 | 0,006±0,003 | 0,004±0,001 | 0,002±0,001 | *0,279* | *0,317* |
| **Body Speed (cm/s)** | 36,275±5,619 | 35,296±5,597 | 39,803±2,571 | 38,306±2,252 | *0,787* | *0,820* |
| **Body Speed Variation (%)** | 19,484±2,633 | 19,599±2,386 | 18,686±2,399 | 20,481±2,643 | *0,935* | *0,548* |
